# A method for generating user-defined circular single-stranded DNA from plasmid DNA using Golden Gate intramolecular ligation

**DOI:** 10.1101/2022.11.21.517425

**Authors:** Isabell K. Strawn, Paul J. Steiner, Matilda S. Newton, Timothy A. Whitehead

## Abstract

Construction of user-defined long circular single stranded DNA (cssDNA) and linear single stranded DNA (lssDNA) is important for various biotechnological applications. Many current methods for synthesis of these ssDNA molecules do not scale to multi-kilobase constructs. Here we present a robust methodology for generating user-defined cssDNA employing Golden Gate assembly, a nickase, and exonuclease degradation. Our technique is demonstrated for three plasmids with insert sizes ranging from 2.1-3.4 kb, requires no specialized equipment, and can be accomplished in five hours with a yield of 33-43 % of the theoretical. To produce lssDNA, we evaluated different CRISPR-Cas9 cleavage conditions and report a 52 ± 8% cleavage efficiency of cssDNA. Our method presented here can make ssDNA readily available to biotechnology researchers.

## Introduction

Single-stranded DNA (ssDNA) is increasingly becoming important for novel techniques and advancements in biotechnology, synthetic biology, and biomaterial research. Long ssDNA molecules have a wide range of potential applications including genome engineering and recombineering, DNA storage, DNA origami, and gene therapy [1]. There is a need for an accessible method of synthesis of user-defined ssDNA that is cost-effective, simple, robust, and scalable.

One major application of ssDNA is its use as a template for genome engineering applications. RNA-programmable gene targeting nucleases like CRISPR-Cas9 have transformed genome engineering in animals [2]. Site-specific modification of genes and full allelic replacement can occur through double stranded breaks (DSB) at defined loci, followed by Homology Directed Repair (HDR) with a donor nucleic acid molecule containing homology arms. However, inserting long contiguous DNA is technically challenging. A grand challenge in genome editing is the highly efficient site-specific insertion, or replacement, of larger DNA cassettes into mammalian genomes. Solving this challenge is important for several fields spanning discovery biology and biotechnology: (i.) generation of transgenic animal models including conditional knockout mice and mice with inducible transcriptional activators [3]–[7]; (ii.) high efficiency insertion of transgenes in cell lines at site-specific locations [8]; (iii.) development of diverse antibody libraries using mammalian cell display [9]; and (iv.) insertion of chimeric antigen receptors in T cells for immunotherapy [10], [11]. More specifically, *Easi-*CRISPR [7], [12] provides a partial solution to the challenge of efficient DNA cassette insertion in mammalian genome engineering using a long, linear single-stranded DNA (lssDNA) donor as opposed to double stranded DNA [13]–[17] or overlapping short ssDNA molecules.

Current ssDNA synthesis strategies can be complex, expensive, time consuming, prone to high error rates, or rely on specialized equipment, and many of the current methods do not scale well over 2 kilobases in length. Commercial vendors such as IDT offer chemical synthesis of long ssDNA. The traditional method of chemical synthesis involves using phosphoramidite chemistry on solid surfaces [18]. However, chemical synthesis is oftentimes limited to smaller ssDNA fragments. Veneziano et al showed that gene-length lssDNA donors could be produced by enzymatic synthesis techniques such as asymmetric PCR (aPCR) [19]. aPCR involves two amplification primers with one in excess to produce the desired ssDNA, but it can require extensive experimentation to optimize yield of the ssDNA and the technique can frequently result in the creation of unwanted byproducts and nonseparated DNA strands [1], [19]. Alternatively, rolling circle amplification can generate long repeating ssDNA using an isothermal polymerase reaction to continuously anneal deoxynucleotide triphosphates to a DNA primer [20], [21]. While this technique is robust, the reaction time can be long, the amplification efficiency is low, and long plasmid-length DNA encoded includes unwanted bacterial control elements like bacterial origins of replications [1]. There are a wide range of additional techniques for ssDNA synthesis such as transcription and reverse transcription, separation of ssDNA from dsDNA, and a primer exchange reaction [1].

We aimed to develop a robust method to generate long, single-stranded DNA from bacterial-prepped plasmid DNA [7]. The method presented here is cost-effective, simple, and uses readily available materials to generate ssDNA in a short time. A series of enzymatic reactions including a Golden Gate assembly, digestion with nickase, and exonuclease degradation allows for synthesis of a user-defined long, circular ssDNA. By having the user-defined insert of interest (IOI) contained in a plasmid initially and using these straightforward steps, this process can be scaled up for larger microgram ssDNA quantities for downstream applications.

## Results

Our initial overall goal was to develop a robust, efficient method that can prepare long circular and linear ssDNA containing a user defined insert of interest (IOI) from plasmid dsDNA. Our constraints for any new method were that it be easily scalable for longer IOI segments, require no specialized equipment or reagents, be robust towards different insert sequences, have high yield of ssDNA products, remove the plasmid backbone section from the final ssDNA product, and be a single-day procedure.

We first evaluated a method where the first step involved preparing ssDNA of the entire plasmid using a nickase and exonuclease digestion. Then, ‘masking oligonucleotides’ (primers that can protect the ssDNA IOI by creating a dsDNA barrier against selective exonuclease digestion) adjacent to the IOI could be annealed and extended by Phusion polymerase to form dsDNA product around the plasmid backbone. The IOI ssDNA would be preserved by masking primers containing a dideoxycytosine modification that would disrupt Phusion polymerase action. Finally, degradation with a dsDNA-specific nuclease would result in liberation of the linear ssDNA IOI. We evaluated several different dsDNAses and found that Evrogen dsDNAse was most selective between ssDNA and dsDNA. Initial tests did not result in reliable reproduction of linear ssDNA (data not shown). We hypothesized that polymerase read-through hindered ssDNA production, and so next attempted the masking reaction with 5-bromo-deoxyuridine (5BU)-modified primers. This modification allowed for the flanking primer to anneal irreversibly to the ssDNA plasmid via UV-induced crosslinking between DNA strands. While this technique showed some specific degradation, again the results were inconsistent or not reproducible.

Based on these results, we attempted an alternative strategy involving a Golden Gate intramolecular ligation of the insert (**Figure 1a)**. In the first step, Golden Gate assembly generates, among other products, a dsDNA plasmid that contains only the insert through intramolecular ligation. In the second step, a cleanup step involving restriction digestion and exonuclease treatment removes unwanted byproducts such as the linearized DNA and circular plasmid backbone. In the third step, we generate IOI-specific circular ssDNA (cssDNA) using a nickase and exonuclease digestion; this process leaves a scar site (< 7 nt at either the 5′ or 3 ′ end) introduced to accommodate the nicking process. In the final step, cssDNA is linearized at the scar site or any selected region using an sgRNA-Cas9 construct, resulting in user-defined linear ssDNA.

**Figure 1:**
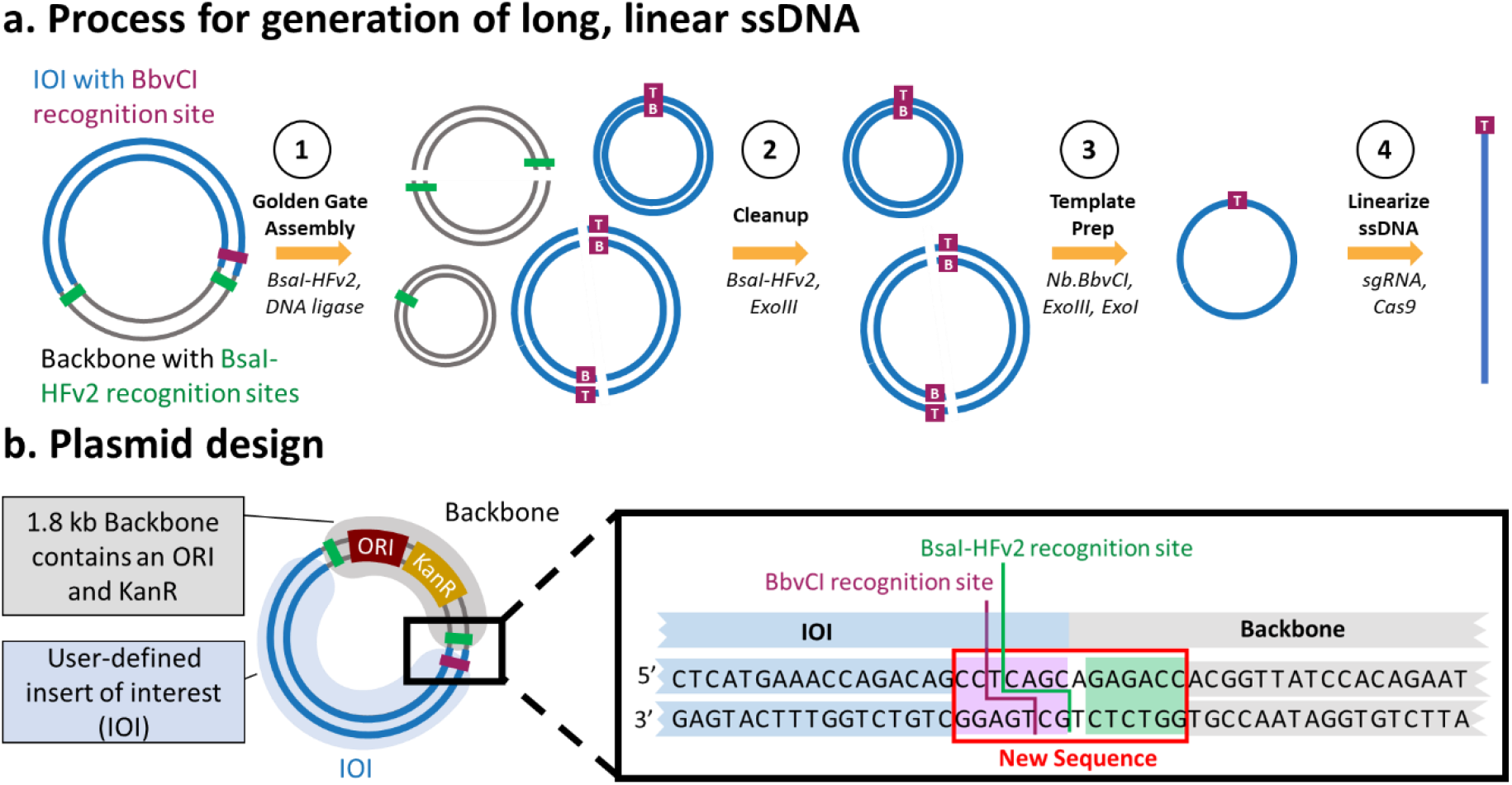
Protocol overview for generating custom lssDNA from plasmid DNA. **a. Process for generation of long, linear ssDNA.** Starting from a plasmid containing the insert of interest (IOI; blue), Golden Gate assembly is used to generate an intramolecular ligation product of the IOI. Undesired reaction products like backbone (gray) and concatemer plasmids of the insert are shown (Step 1). The two strands of the plasmid are labelled as T (top) and B (bottom), referencing the preference of the enzyme BbvCI. A cleanup step using BsaI-HFv2 and Exonuclease III removes undesired backbone reaction products (step 2). Then, circular ssDNA can be created from dsDNA using a nickase followed by exonuclease degradation (step 3). Finally, the cssDNA is linearized by Cas9 cleavage at the BbvCI recognition site, leaving a maximum of a 7 nt 5’ or 3’ scar (step 4). **b. Plasmid design**. The IOI (blue) is ligated into a backbone (grey) containing a high copy number ORI, a KanR resistance cassette, a BbvCI recognition site, and flanking BsaI-HFv2 recognition sites. The inset shows the sequence design of the adjacent BbvCI and BsaI-HFv2 recognition sites.

A plasmid compatible with this process would include (i.) a minimal backbone enabling selection and replication in *E. coli;* (ii.) a BsaI restriction site adjacent to the insert for intramolecular ligation in the Golden Gate assembly; and (iii.) a flanking BbvCI restriction site for nickase digestion (**Figure 1b**). The insert cannot contain any BsaI recognition sequence, and any BbvCI restriction sites must occur on the same sense strand of DNA. Plasmid pIS001 (5161 bp), a test plasmid compatible with the strategy, was prepared by Gibson assembly [22] of the pYTK084 backbone [23] containing a high copy number ORI and a kanamycin resistance cassette with the 3379bp insert ROSA-TRE-C16V5-G-pA. This insert contains 5’ and 3’ homology arms of mouse ROSA26 locus, TRE Tight, CEACAM16 coding sequence V5 tag, EGFP, and BGH polyA.

To optimize the process of ssDNA synthesis, we individually tested each of the four steps for functionality, identified byproducts, and tested variable conditions on distinct steps. Optimization for steps one (Golden Gate intramolecular ligation) and two (reaction clean-up) were performed on pIS001. For step three (formation of cssDNA) we used an existing nicking/digesting protocol that is part of the nicking mutagenesis technique previously developed by our lab [24] which was performed without optimization. Finally, step four (lssDNA generation) was performed on the bacteriophage DNA M13mp18 as both dsDNA and cssDNA are commercially available.

We first confirmed the Golden Gate intramolecular ligation of the ROSA-TRE-C16V5-G-pA insert in a 20 µL reaction volume performed in 1X T4 DNA ligase buffer containing 1 µg pIS001, BsaI-HFv2 (1 µL of a 20 U/µL stock), and T4 DNA ligase. The reaction was subjected to 30 cycles of 37°C incubation for one minute followed by 16°C for one minute, and then a single 60°C incubation for 5 minutes. Digestion with only BsaI-HFv2 resulted in two distinct bands at the approximate sizes of backbone (1.8 kb) and insert (3.4 kb) (**Fig 2a**). When T4 DNA ligase was added to the reaction mixture, it generated additional bands (**Fig 2a**) presumably corresponding to monomers and concatemers of different insert and backbone ligation products. We hypothesized that the band at circa 2 kb corresponded to the desired intramolecular ligation product of the insert and sought to optimize its formation by testing different reaction volumes (20 mL, 100 mL) and BsaI-HFv2 amounts (1, 3 µL of a 20 U/µL stock). We observed a lower abundance of the linearized insert at the higher BsaI-HFv2 amount of 3 µL (**Figure 2a**). With respect to increased reaction volume, we hypothesized that decreasing the overall concentration of linearized IOI fragments would result in fewer concatemers and more of the putative desired intramolecular ligated insert. Our experiments supported this hypothesis: we found that 100 µL reaction volume resulted in higher concentration of circularized insert monomer as judged by gel densitometry (**Figure 2a**). Increasing the reaction volume to 200 µL did not appreciably increase the concentration of monomer but it did qualitatively decrease the formation of higher order concatemers (**Figure 2b**). Thus, a reaction volume of 200 µL and a BsaI-HFv2 amount of 60 U (3 µL of 20 U/µL) was used for all further experiments. One disadvantage of the larger reaction volume is that the downstream steps work in a smaller reaction volume, and so an additional DNA concentration column clean-up is necessary between steps one and two.

**Figure 2:**
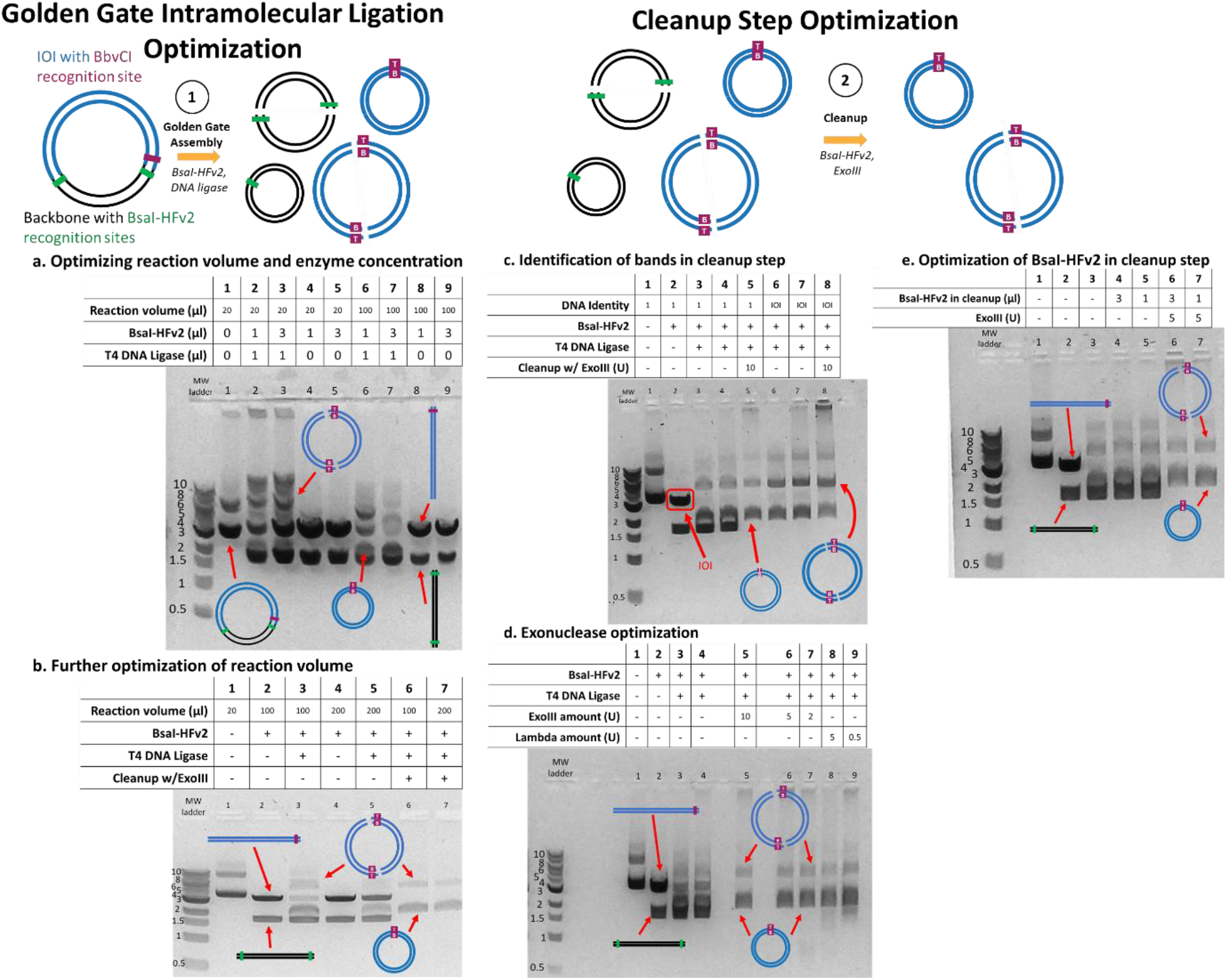
Optimization of the intramolecular ligation by Golden Gate Assembly and of the cleanup step. All lanes showing reaction intermediates contain a starting amount of 1 µg of pIS001 dsDNA. **a**. Optimization of reaction volumes of 20 µl and 100 µl and two different BsaI-HFv2 concentrations (1 µl or 3 µl of a 20 U/µl stock). Higher BsaI-HFv2 and the larger volume resulted in less linearized IOI, and lower concatemers than other conditions. **b**. A larger reaction volume of 200 µl resulted in similar formation of monomeric linear plasmid and formation of concatemers when 3 µl BsaI-HFv2 was included. Lanes 6 and 7 show removal of linear DNA using ExoIII. **c**. Identification of bands in cleanup step. The aim of this experiment was to confirm the identity of the upper bands that appeared in the cleanup step. We first performed BsaI-HFv2 digestion of pIS001 and gel extracted the IOI (red box, lane 2). Next, we performed the Golden Gate intramolecular ligation and cleanup steps on pIS001 (lanes 3-5) and the linearized IOI (lanes 6-8). Based on similar migration of the IOI specific lanes, we conclude that the upper bands were concatemers formed during Golden Gate Assembly. **d**. ExoIII and lambda exonucleases were tested for functionality in the cleanup step at different dilutions using Golden Gate assembly conditions optimized in panel b. Both exonucleases were able to remove linearized DNA while not appreciably degrading circularized IOI at all concentrations evaluated. **e**. Optimization of BsaI-HFv2 in cleanup step (1 µl or 3 µl of a 20 U/µl stock).

Following column clean-up, we tested exonuclease digestion using a 1:10 dilution of Exonuclease III (1 µL of a 1:10 dilution made in 1X CutSmart buffer from 100 U/µL stock ExoIII in a 20 µL total reaction volume). This reaction was sufficient to remove the linear backbone (**Figure 2c**), resulting only in intramolecular ligation products that we hypothesize to be monomers and concatemers of insert sequences. To test this hypothesis, we performed a gel extraction of the linearized insert following BsaI-HFv2 digestion of pIS001 (**Figure 2c**). Next, we performed the intramolecular Golden Gate assembly step on the gel extracted linearized insert, resulting in the same band sizes and distributions as seen by Golden Gate assembly and exonuclease cleanup of the full pIS001 plasmid (**Figure 2c**). From these experiments, we conclude that the bands observed correspond to the desired circularized insert and undesired concatemer side products.

Further optimization was performed on the cleanup step by evaluating different concentrations of exonucleases (Lambda, ExoIII) able to degrade linear dsDNA with a variety of end geometries. Both exonucleases worked sufficiently in cleaning up any remaining linear DNA and any backbone concatemers, leaving only the circular dsDNA plasmid of the insert (**Figure 2d**). In the optimized procedure we used 5 U of ExoIII (1 µL of a 1:20 dilution in 1X CutSmart Buffer of 100 U /µL in a 20 µL reaction volume), although we note that 2 U (1:50 dilution) ExoIII does not show qualitative differences in circularized insert yield and presence of linear contaminants. Two different concentrations of BsaI-HFv2 were tested in the cleanup step (1, 3 µL of a 20 U/µL stock), with the lower concentration sufficient to remove any circularized backbone as judged by gel densitometry (**Figure 2e**). Finally, we used analytical gel densitometry to estimate the mass percentage of monomers to the total DNA mass observed on the gels in the cleanup step. The average percent mass of monomers (over total mass including concatemers) was calculated over four independent gel runs using pIS001. Based on gel densitometry, 75 ± 7% (n = 4, error represents 1 s.d.) of the DNA visible by gel electrophoreses after the cleanup step is the desired dsDNA monomer IOI product.

Given our optimization of steps one and two, and previous optimization of cssDNA production (step 3) [24], we applied the complete protocol to pIS001 to produce cssDNA for the ROSA-TRE-C16V5-G-pA insert (**Figure 3**). We digested reaction products from step two with either Nt.BbvCI or Nb.BbvCI in order to demonstrate that cssDNA could be produced in either the sense or antisense direction (**Figure 3**). This overall procedure from a 1 µg plasmid DNA input took a minimum of 4.5 h: 1.5 h for the Golden Gate intramolecular ligation, 0.25 h for column clean-up, 1.25 h for the cleanup step, and 1.5 h for the template preparation step.

**Figure 3.**
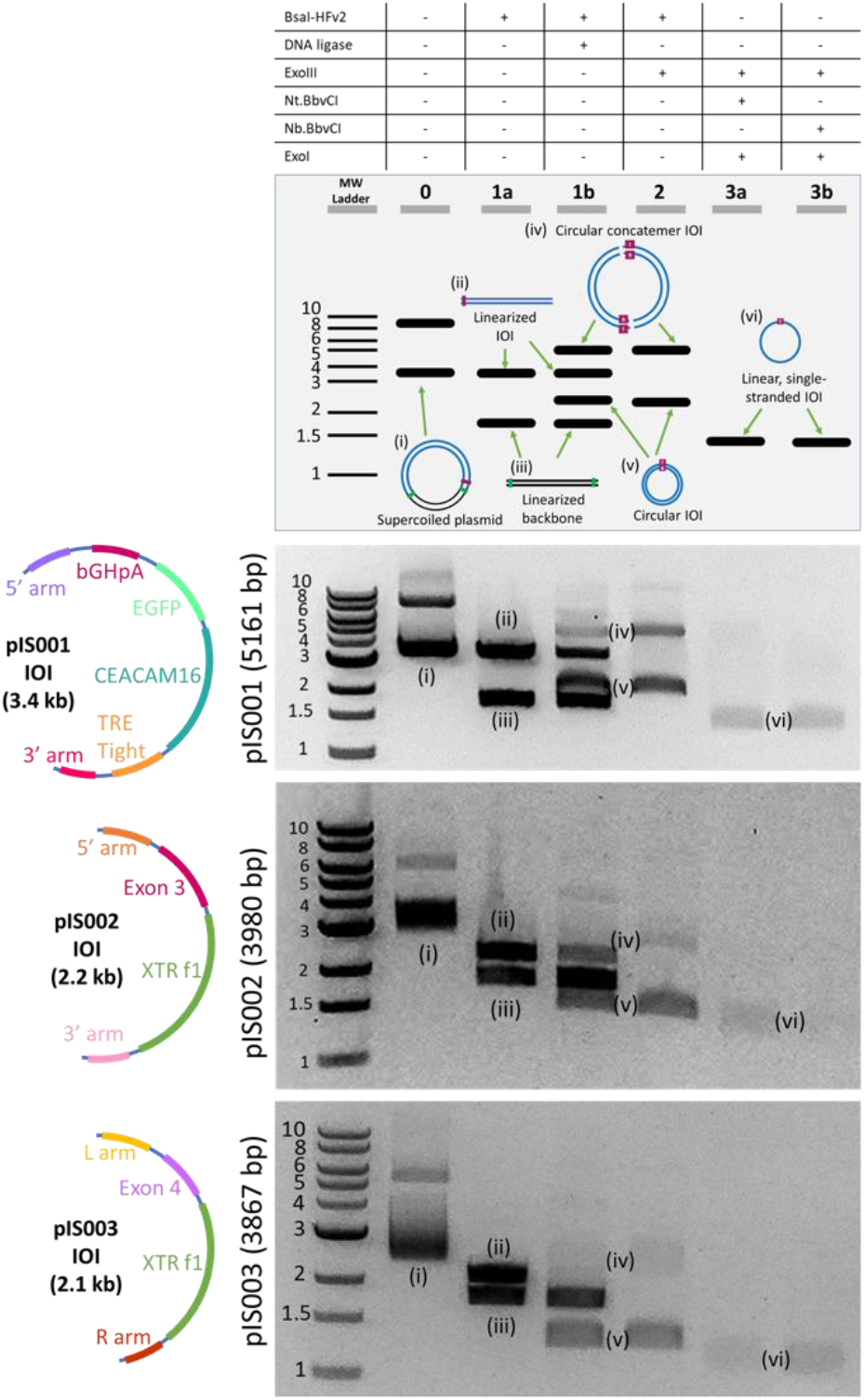
cssDNA productions for three separate inserts. The optimized protocol was tested on the three different plasmids that were constructed (pIS001, pIS002, and pIS003), with inserts shown in the first column. The results of each step of the process for 1 µg of pIS001 (5161 bp), pIS002 (3980 bp), and pIS003 (3867 bp) are shown. Lanes correspond to the steps in the protocol listed in Figure 1. Lane 0: undigested plasmid DNA containing supercoiled and linearized plasmids. Lane 1a: plasmid is digested with BsaI-HFv2. Lane 1b: Golden Gate Assembly including BsaI-HFv2 and DNA ligase. Lane 2: Cleanup step following Golden Gate assembly. Lanes 3a-b: cssDNA is visible when either the Nt.BbvCI (3a) or Nb.BbvCI (3b) nickase is used for template prep. ssDNA bands do not appear as brightly as dsDNA in part because the SYBR-Safe dye used has a lower quantum yield when bound to ssDNA than dsDNA at the wavelength used for imaging.

To demonstrate that this method robustly produces cssDNA for distinct insert sequences, we prepared two additional plasmids: pIS002 (3980 bp) contains a 5’ arm, CEACAM16 Exon3, XTR cassette [25], and 3’ arm, and pIS003 (3867 bp) contains a 5’ arm, CEACAM16Exon4, XTRf1, and 3’ arm. pIS002 also contained an additional BbvCI site, and the nicking step was performed with the additional BbvCI site in the same direction as the existing site so that this did not impact the results. For both plasmids, our method could successfully produce insert cssDNA in either the sense or antisense configurations (**Figure 3**). The cssDNA production step appeared to remove higher order concatemers, which we attribute to incomplete ligation of the concatemers by the T4 DNA ligase. For example, a plasmid dimer has four nicks to seal, whereas a monomer only has two, and any nicks on the strand complementary to the nickase would result in full degradation of dsDNA. Finally, we performed three technical replicates of our protocol for both pIS001 and pIS002 and evaluated the cssDNA yield using a Qubit ssDNA Assay Kit. For an input of 1 mg of plasmid dsDNA, we observed 109 and 120 ng of cssDNA for pIS001 and pIS002, respectively, resulting in a theoretical yield of 33±2% (pIS001) to 43±2% (pIS002) (**Table 1;** error represents 1 s.d.). Note that the quantification assay required an additional concentration step that may reduce yield by up to 30%, so the true yield is likely marginally higher than what is reported.

**Table 1.** cssDNA yield calculations for the optimized cssDNA protocol. Error bars report 1 s.d., n=3 technical replicates.

| **Plasmid** | **Yield (ng/1 μg input DNA)** | **Maximum Theoretical Yield (ng/ 1 μg input DNA)** | **Yield (% theoretical)** |
| --- | --- | --- | --- |
| pIS001 | 109±7 | 327 | 33±2 |
| pIS002 | 120±6 | 276 | 43±2 |

For producing linear ssDNA from cssDNA, we sought a commercially available endonuclease capable of site-specific cleavage of ssDNA. While CRISPR-Cas9 functions predominantly on dsDNA, Sternberg and Ma reported that diverse Cas9 proteins could cleave ssDNA, albeit at a reduced yield [26], [27]. In particular, they reported that the common and commercially available *Spy*Cas9 cleaves ssDNA at approximately 50% yield on short ssDNA constructs. To evaluate *Spy*Cas9, we used M13mp18 (7249 nts) circular ssDNA and dsDNA, as both are commercially available. We produced a sgRNA specific for M13mp18 and first addressed whether the sgRNA-*Spy*Cas9 complex could cleave M13mp18 dsDNA (**Figure 4a**). We found cleavage only when both sgRNA and *Spy*Cas9 were present in the reaction mixture, confirming sgRNA functionality. Next, we tested sgRNA-*Spy*Cas9 functionality on circular ssDNA (**Figure 4b**). We observed appreciable cleavage at all incubation temperatures tested (**Figure 4b**) but no improvement at any temperature over the approximately 50% yield similar to the observations by Sternberg (2014) and Ma (2015). Through gel densitometry, we estimated a 52 ± 8% linearization efficiency (n = 3, error represents 1 s.d.). We also tested various Cas9:sgRNA:DNA molar ratios (5:5:1; 10:10:1; data not shown), but the optimization results did not yield any further improvements in lssDNA production.

**Figure 4.**
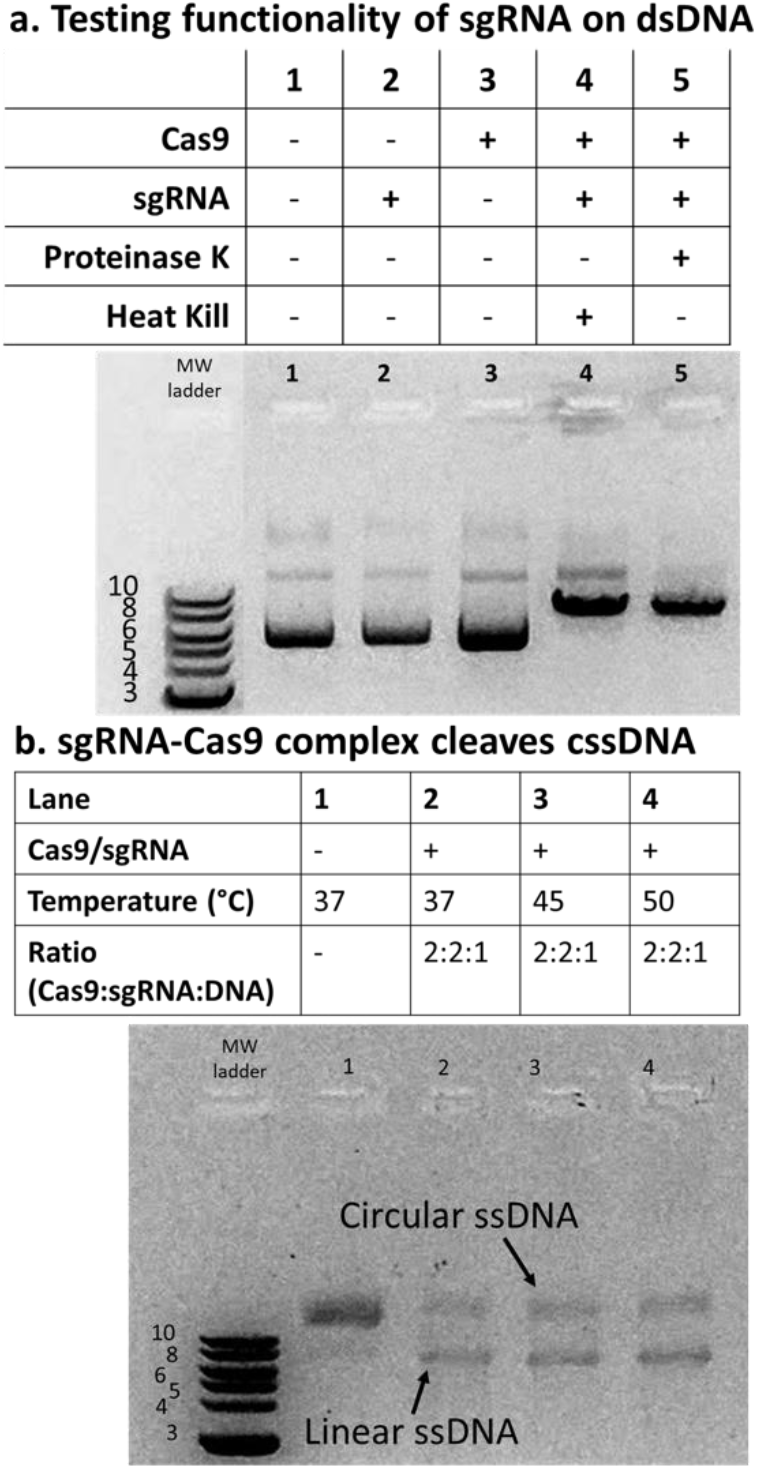
sgRNA-*Spy*Cas9 complex cleaves cssDNA to produce lssDNA. The linearization of the DNA using sgRNA-*Spy*Cas9 complexes was tested using 1 µg of M13mp19 dsDNA or ssDNA. **a**. Testing functionality of sgRNA. *Spy*Cas9 and designed sgRNA could cleave dsDNA only when both were included in the reaction (lanes 4-5). Proteinase K or 60°C incubation was used to dissociate the RNP from DNA. **b**. sgRNA-*Spy*Cas9 complex cleaves cssDNA with moderate efficiency. ssDNA cleavage using the sgRNA-Cas9 complex was successful but had poor yield. In order to properly visualize and differentiate linear ssDNA from circular ssDNA, the DNA was run on a 1.5% w/v gel with the addition of 60% v/v formamide followed by heating at 70°C for 5 minutes and then on ice for 5 minutes (to help denature the ssDNA and result in clearer bands). Variable temperatures did not increase yield.

## Discussion & Conclusion

Our method presented here can be used to prepare cssDNA at a theoretical yield of up to 43±2% using commercially available reagents and in less than 5 h. The robustness of the method was demonstrated using three differently sized inserts 2.1-3.4 kb in length. Based on these results, we hypothesize, but did not demonstrate, that we could accommodate larger nucleotide inserts as well. Follow-up work could test the upper size limits of this protocol. One potential limitation of our cssDNA protocol for downstream applications like genome editing is the introduction of a 7 nt scar site on the insert. In theory, this scar site can be removed if the insert already contains an endogenous BbvCI recognition site in an opportune location. It remains to be demonstrated whether this 7 nt scar impacts insertion efficiencies using long 5’ and 3’ homology arms.

We were unable to demonstrate a high efficiency protocol for preparation of lssDNA using an endonuclease capable of cleaving ssDNA. Using commercially available *Spy*Cas9 with a sgRNA specific for bacteriophage ssDNA, we were able to achieve a cleavage efficiency of approximately 50%, and neither higher temperature nor higher molar ratios of *Spy*Cas9:sgRNA compared with ssDNA resulted in appreciable increases in cleavage efficiency. It is known that sgRNA-SpyCas9 complex can bind to ssDNA, but the efficiency of the cleavage step is dramatically reduced relative to dsDNA [26]. We calculated with gel densitometry that 52 ± 8% of the bacteriophage M13mp18 cssDNA was cleaved to linear ssDNA. Sternberg et al., showed that other Cas9 proteins, like *Cdi*Cas9, could cleave ssDNA at higher yields. *Cdi*Cas9 and other high-yield ssDNA cutting Cas9 were not explored in depth here due to the criterion of the reagents being readily available; this should be a prime focus in future work. An alternative future direction is to test rare type II restriction enzymes that have been reported to cleave ssDNA [28].

The ssDNA produced in this study can be used for a variety of biotechnology applications. One major application of ssDNA is for inserting long contiguous DNA into animal models using *Easi-*CRISPR. *Easi-*CRISPR uses long ssDNA donors injected with crRNA + tracrRNA + Cas9 ribonucleoprotein (ctRNP) complexes into mouse zygotes to efficiently perform genome engineering. SsDNA can also be implemented in other variations of templated homology directed repair (HDR) for genome editing because ssDNA increases the likelihood of HDR with reduced cellular toxicity [11], [19], [29]–[31]. Another potential application includes DNA origami which is limited by the availability of diverse single-stranded DNA scaffolds that are required for the method. Self-assembling DNA nanostructures can be designed and constructed using DNA origami-folding, which involves using single-stranded DNA to direct the folding of a long ssDNA scaffold [32].

In conclusion, we have demonstrated that user-defined circular ssDNA can be easily and cost-effectively generated using widely available reagents and basic lab techniques. We were unable to demonstrate conversion of cssDNA to linear ssDNA at a higher efficiency, but there is room for further optimization for this step and it is possible that with further experimentation, a high efficiency linear ssDNA can be synthesized using this method. Regardless, the method proposed here for cssDNA synthesis could be easily scaled up as needed to produce larger quantities of ssDNA at a relatively low cost and promises to make ssDNA creation a more affordable and accessible process for researchers.

## Methods

### Reagents

All DNA modification enzymes, buffers, and reagents were sourced from New England Biolabs (NEB) unless otherwise noted. All primers and oligos were ordered from Integrated DNA Technologies (IDT).

### Plasmid Construction

All primer sequences along with any oligos used in this work are listed in **Supplementary Table 1**, and all plasmids are listed in **Supplementary Table 2**. Plasmids C16Ex2, C16Ex3, and C16Ex4 were custom synthesized by Azenta life sciences. pIS001 was constructed with Gibson Assembly from plasmid C16Ex2 containing the ROSA-TRE-C16V5-G-pA insert and the pYTK084 backbone [23]. Primers p001 and p002 were used to amplify insert and primers p003 and p004 were used to amplify the backbone, and fragments assembled by Gibson Assembly using NEBuilder® HiFi DNA Assembly Master Mix (NEB; catalog # E2621L). Plasmids pIS002 and pIS003 were also constructed using Gibson Assembly similarly to pIS001, except the inserts were isolated from plasmids C16Ex3 and C16Ex4, respectively, using a restriction digest with BsaI-HFv2 (NEB; catalog # R3733L) in 1X rCutSmart Buffer and subsequent gel extraction.

### Intramolecular Golden Gate Assembly and Cleanup

All Golden Gate Assembly reactions started with 1 µg of the constructed plasmid (pIS001, pIS002, or pIS003) and BsaI-HFv2 (20 or 60 U from 20 U/µL) (NEB; catalog # R3733L), 400 U of T4 DNA ligase (1 µL of 400 U/µL) (NEB; catalog # M0202L), and 1X T4 DNA ligase buffer (NEB; catalog # B0202S). During optimization, reactions were performed either in 20, 100, or 200 µL reaction volumes. Reaction mixture was placed in a Mastercycler X50s thermocycler for 30 cycles of 37°C for 1 minute followed by 16°C for 1 minute, followed by a 60°C hold for 5 minutes. Following the Golden Gate Assembly step, the reaction was cleaned and concentrated with a Monarch PCR & DNA Cleanup Kit (NEB catalog # T1030L) and eluted in either 10 or 12 µL of nuclease free H2O. This elution product was then incubated with BsaI-HFv2 (20 or 60 U from 20 U/µL), 1X rCutSmart buffer (10X), nuclease free H20, and an exonuclease (1 µL of either Lambda or Exonuclease III) in a 20 µL cleanup reaction. During optimization, we tested different amounts of Exonuclease III (NEB; catalog # M0206L) (2, 5, 10 U) made by dilution of the stock 100 U /µL using 1X rCutSmart Buffer, or Lambda (NEB; catalog # M0262L) (0.5,5 U), with the former made by 1:10 dilution of the 5 U/µL stock. This reaction was incubated in a thermocycler at 37°C for 30 minutes followed by 16°C for 20 minutes.

### Template Prep Step for Production of cssDNA

All template preparation reactions occurred in 25 µL and were comprised of the 20 µL cleanup step, 1.5 µL of nuclease free water, 10 U of either Nb.BbvCI (NEB; catalog # R0631L) or Nt.BbvCI (NEB; catalog # R0632L (1 µL of 10 units/µL), 10 U of Exonuclease III (1 µL of a 1:10 dilution in 1X rCutSmart Buffer of 100 U/µL stock), 20 U of Exonuclease I (1µL of 20 U/µL) (NEB; catalog # M0293L), and 0.5 µL of 10X rCutSmart Buffer. This reaction was placed in the thermocycler at 37°C for 60 minutes followed by 16°C for 20 minutes. The results were column purified with the Monarch PCR & DNA Cleanup Kit and eluted in either 10 or 12 µL of nuclease free water.

### sgRNA Production and Cas9-sgRNA Cleavage Experiments

To make the sgRNA, the EnGen sgRNA Synthesis Kit, *S. pyogenes* (NEB; catalog # E3322V) was used with a DNA primer as a template (**Supplementary Table 1**). The linearization of the cssDNA was performed in three steps: a pre-cycle, a cutting step, and inactivation of Cas9. In the pre-cycle step, 3 µL of 300 nM sgRNA diluted in NFH_2_O, 1 µL of 1 µM Cas9 Nuclease (diluted from 20 µM stock in 1X NEBuffer r3.1) (NEB; catalog # M0386T), and 1X NEBuffer 3.1 (NEB; catalog # B6003S) were combined in a 20 µL reaction volume, and the solution was placed in the thermocycler at 25°C for 10 minutes. In the digestion step, 1 µg of M13mp18 ssDNA (NEB; catalog # N4040S) was added to the pre-cycle results for a total reaction volume of 30 µL. This reaction was placed in the thermocycler for 60 minutes at 37°C. For the inactivation step, we added 0.8 U of Proteinase K (1 µL of 0.8 U/ µL) and incubated at room temperature for 10 minutes, or alternatively tested a heat inactivation of 65°C for 15 minutes.

### Gel Imaging and Reaction Conditions

To visualize the results of the procedure, the dsDNA was run on 1% w/v agarose gel in 0.5X Tris-acetate-EDTA (TAE) buffer (made from a 50X comprising 24.2% w/v tris base, 5.71% v/v glacial acetic acid, and 10% v/v 0.5M EDTA, pH8.0) at a range of 100 – 120 V. To run the gels, we used a 1 kb Plus DNA Ladder for Safe Stains (NEB; catalog # N0559S), Gel Loading Dye, Purple (6X) (NEB; catalog # B7024S), and SYBR™ Safe DNA Gel Stain (ThermoFisher Scientific; catalog # S33102). Prior to running on gels, ssDNA for the linearization experiments was denatured by adding 60% v/v formamide solution (formamide with 0.5% v/v EDTA buffered to pH 8.0 with NaOH, filtered through a 0.2µm filter and autoclaved) to the reaction followed by heating at 70°C for 5 minutes and then on ice for 5 minutes [33]. ssDNA was run on a 1.5% w/v agarose gel in 0.5X TAE.

### Quantification of DNA with Gel Densitometry and Fluorescent Assay

To quantify the DNA yield at various steps in the process, ImageJ [34], [35] was used to analyze gels and obtain intensity values from pictures of the bands on the gels. The intensity of the bands is proportional to the mass of the DNA present, and we used this to calculate the percent of a specific DNA product compared to the total DNA mass in the reaction.

The cssDNA yield for pIS001 and pIS002 was calculated using a Quibit ssDNA Assay kit from Invitrogen (ThermoFisher Scientific; catalog # Q10212). cssDNA generated from the template preparation step was column purified using a Monarch PCR & DNA Cleanup Kit and eluted into 11 µL. The cleaned ssDNA samples were prepared according to the ssDNA Assay kit instruction by dilution in 189 µL of Qubit working solution (1:200 dilution of Qubit ssDNA Reagent in Qubit ssDNA buffer) for a total reaction volume of 200 µL. The relative fluorescence units (RFU) of each sample were obtained using a Qubit 3.0 Fluorometer. By comparing the RFU values to a standard curve generated with ssDNA of known concentrations, we quantified the yield nanograms of ssDNA generated. We calculated the maximum theoretical yield as the percentage of IOI in the whole plasmid divided by two to account for the ssDNA product, and calculated the percent yield using these theoretical values.

## Supporting information

Figure 2a gel image

Figure 2b gel image

Figure 2c gel image

Figure 2d gel image

Figure 2e gel image

Figure 3 pIS001 gel image

Figure 3 pIS002 gel image

Figure 3 pIS003 gel image

Figure 4a gel image

Figure 4b gel image

## Acknowledgements

This work was supported by NIH NIGMS under Award Number R21GM129559-01 to T.A.W.

## Data Availability

All original gel image files are given in the supporting information. All Golden Gate compatible plasmids have been deposited at AddGene.

## Notes

### Competing Interest Statement

The authors have declared no competing interest.

