## Supplementary figures and images for "A method for generating user-defined circular single-stranded DNA from plasmid DNA using Golden Gate intramolecular ligation"

### Figure 2a gel image

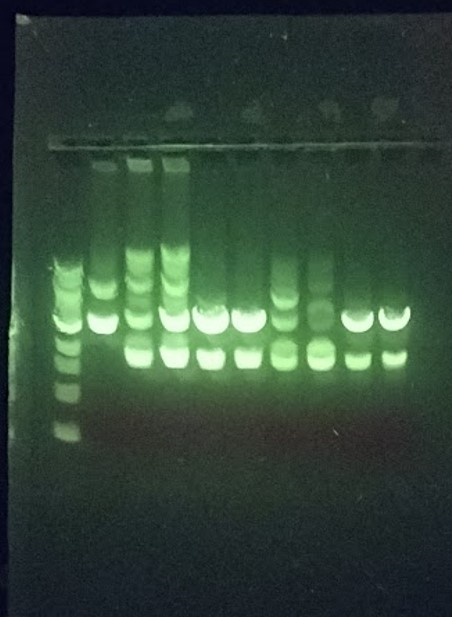

### Figure 2b gel image

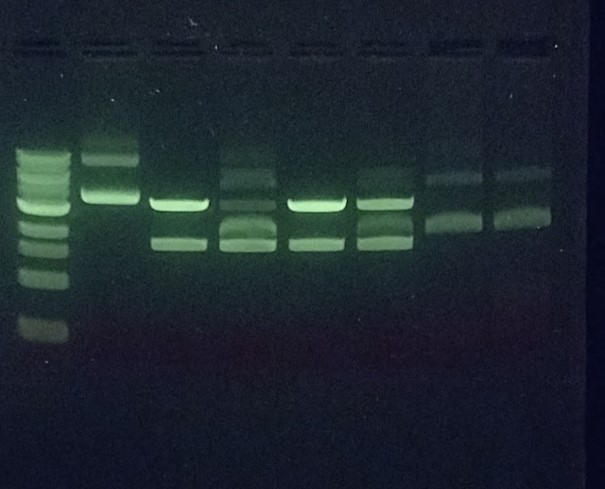

### Figure 2c gel image

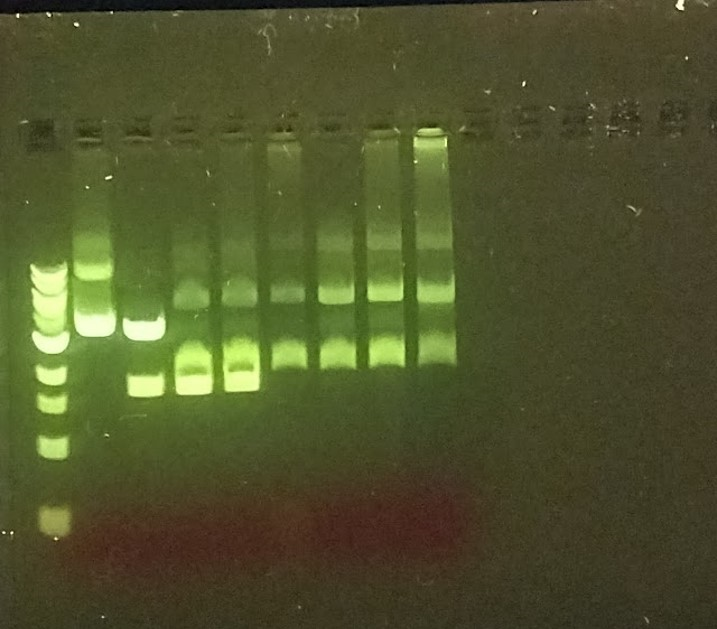

### Figure 2d gel image

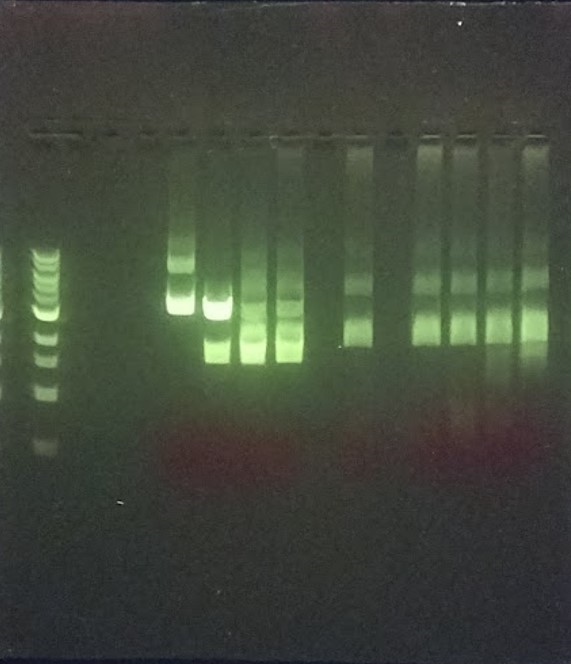

### Figure 2e gel image

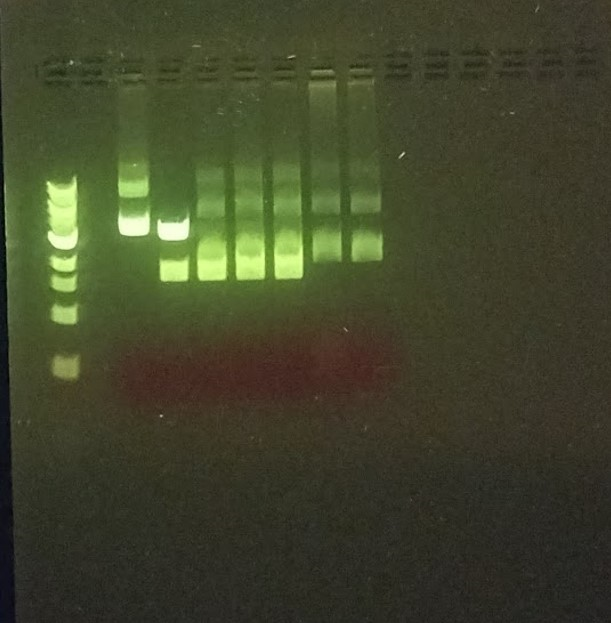

### Figure 3 pIS001 gel image

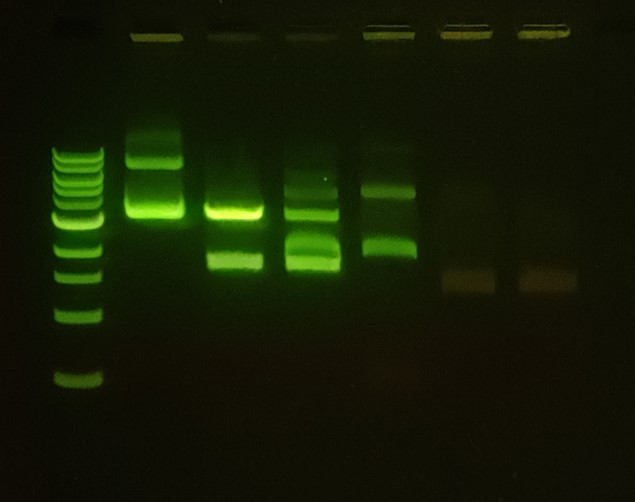

### Figure 3 pIS002 gel image

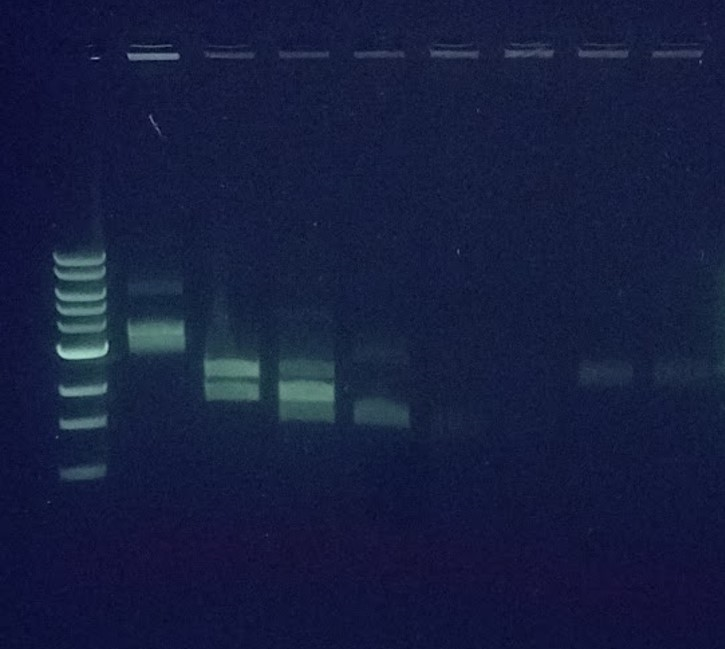

### Figure 3 pIS003 gel image

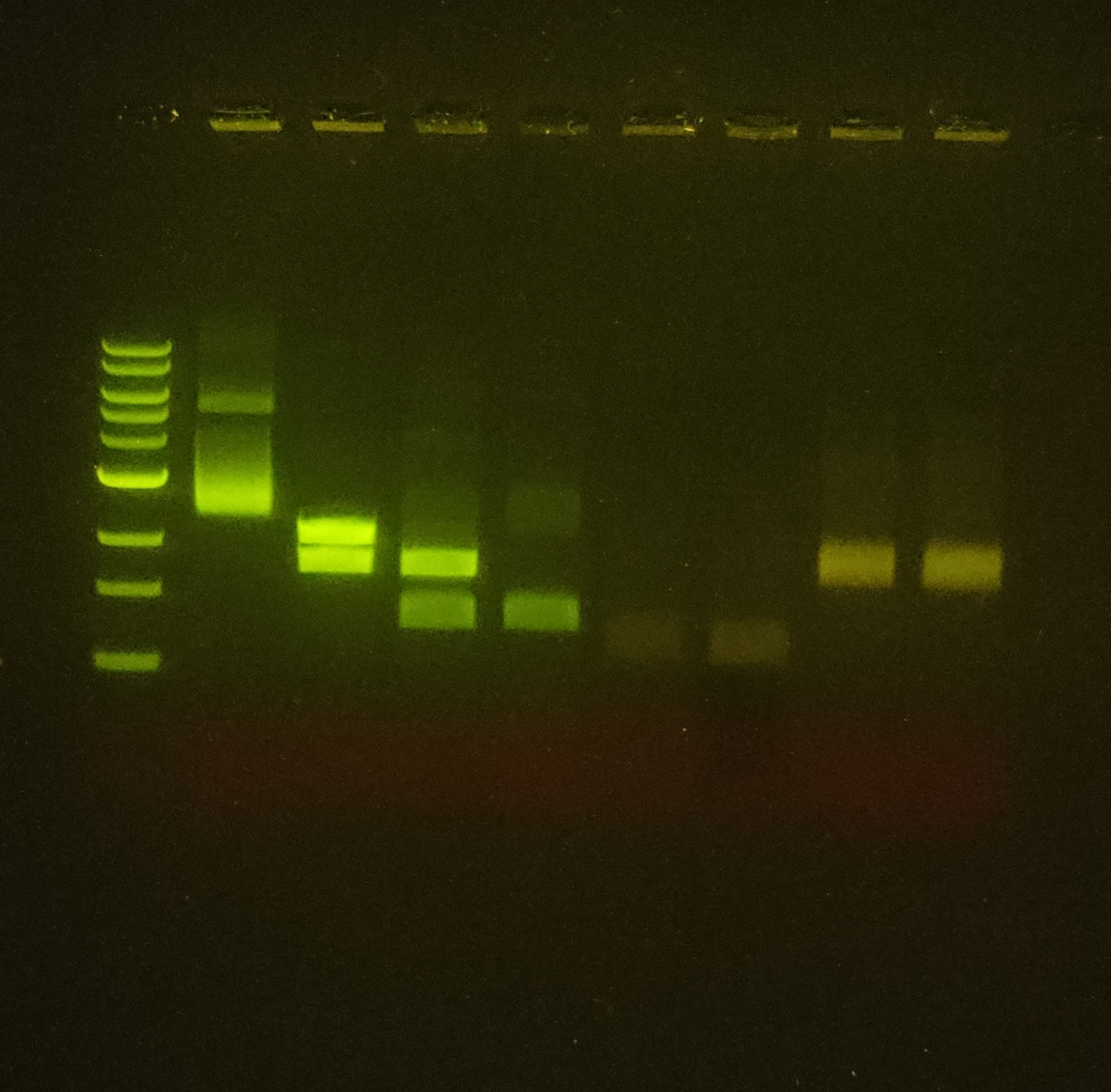

### Figure 4a gel image

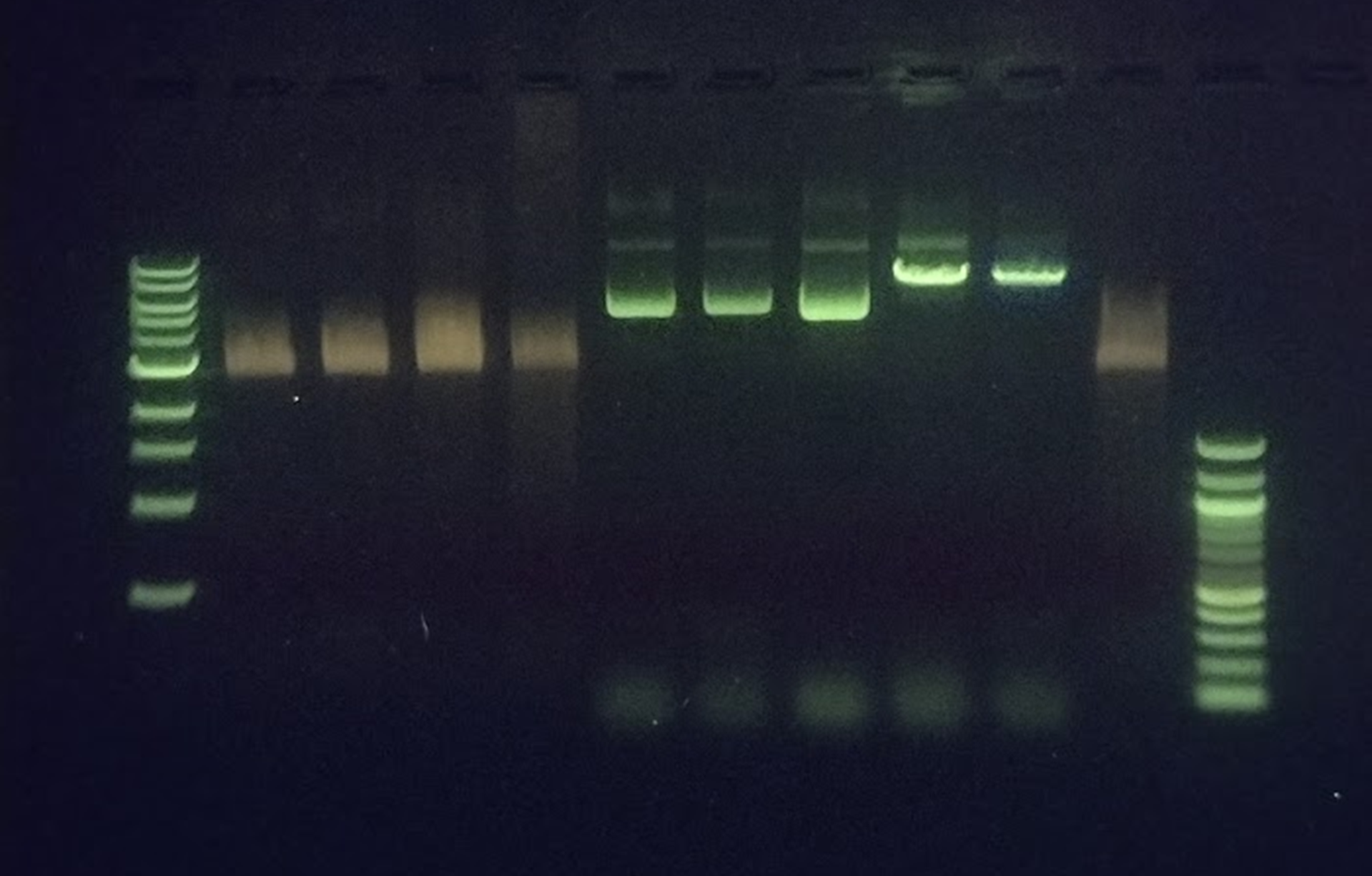

### Figure 4b gel image

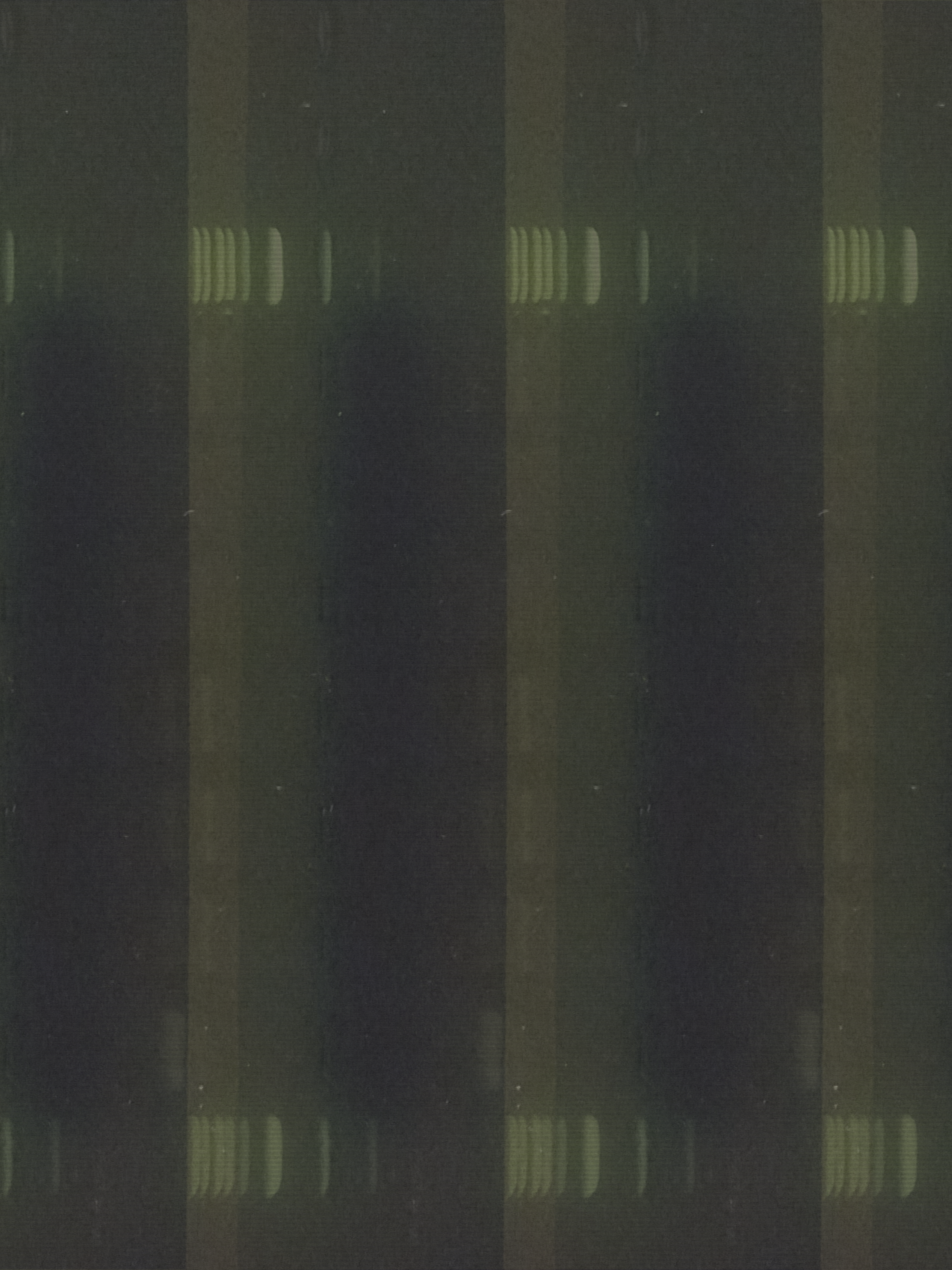
